# Bayesian mixture model for clustering rare-variant effects in human genetic studies

**DOI:** 10.1101/2021.08.03.454967

**Authors:** Guhan Ram Venkataraman, Yosuke Tanigawa, Matti Pirinen, Manuel A. Rivas

## Abstract

Rare-variant aggregate analysis from exome and whole genome sequencing data typically summarizes with a single statistic the signal for a gene or the unit that is being aggregated. However, when doing so, the effect profile within the unit may not be easily characterized across one or multiple phenotypes. Here, we present an approach we call Multiple Rare-variants and Phenotypes Mixture Model (MRPMM), which clusters rare variants into groups based on their effects on the multivariate phenotype and makes statistical inferences about the properties of the underlying mixture of genetic effects. Using summary statistic data from a meta-analysis of exome sequencing data of 184,698 individuals in the UK Biobank across 6 populations, we demonstrate that our mixture model can identify clusters of variants responsible for significantly disparate effects across a multivariate phenotype; we study three lipid and three renal traits separately. The method is able to estimate (1) the proportion of non-null variants, (2) whether variants with the same predicted consequence in one gene behave similarly, (3) whether variants across genes share effect profiles across the multivariate phenotype, and (4) whether different annotations differ in the magnitude of their effects. As rare-variant data and aggregation techniques become more common, this method can be used to ascribe further meaning to association results.

## 1 Introduction

Population-scale sequencing studies are becoming pervasive^1–4^. As a result, analyses considering the joint contribution of rare variants to disease susceptibility and phenotypic variation are also becoming pervasive. Commonly used aggregation approaches for rare-variant association studies include the sequence kernel association test, the burden test, and more-general Bayesian model comparison methods^5–14^. However, aggregation as performed in these methods also tends to lose information within the blocks (typically, genes) specified; that is, they fail to indicate whether certain variants (or certain types of variants) within the same block may have different effects on the multivariate phenotype. In particular, they do not pinpoint which variants are driving the association signal. It is thus critical that we develop methods that can “trace back” variants’ effects, clustering the variants used within the blocks of interest into groups that have distinct per-phenotype aggregate effects while adequately accounting for the uncertainty that rare-variant studies exhibit.

Clustering methods fall into several categories, the most prevalent of which are distance-based methods (such as K-means and self-organizing maps) that are sensitive to the noise that typically plagues biological data^8^. Model-based methods such as mixture models are a simple but elegant alternative that assume that data are generated from multiple source distributions, which are then learned and set as the “clusters”. Algorithms such as Expectation Maximization can estimate latent variables that underlie these clusters. The finite-mixture model, one such model-based method, assumes a finite number of clusters and asserts that this number can be estimated using goodness-of-fit criteria like the Bayesian Information Criterion (BIC) or the Akaike Information Criterion (AIC)^15, 16^.

In this study, we propose to use a Bayesian hierarchical mixture model where a hierarchical structure is introduced to allow the sharing of information among related clusters (in our case, genes) and where the number of clusters are pre-specified. Our approach, the Multiple Rare-variants and Phenotypes Mixture Model (MRPMM), considers several factors when estimating parameters of interest underlying the mixture of effects driving an association signal. We calculate matrices of genetic correlations among phenotypes of interest using both common or null variants and rarer or significant variants. The method also estimates the spread of effects across predicted consequences of the genomic variants (variant annotations), which is a critical aspect of interpreting genetic findings. The annotations represent expected severity of impact on phenotypes (for example, protein-truncating variants, or PTVs^17, 18^, are predicted to truncate the protein product, and are purported to be much more deleterious than other variants). In MRPMM, we use summary statistics (for single-variant single-phenotype GWAS, these are per-variant estimates of marginal univariate effect size and corresponding standard errors). In practice, sharing of individual genotype and phenotype data across groups in large genetic consortia is difficult to achieve due to privacy concerns and consent issues; using summary statistics can bypass these issues while also increasing computational efficiency without reducing accuracy. Insights from Liu et al.^19^ and Cichonska et al.^20^ also suggest that the use of additional summary statistics, like covariance estimates across variants and studies, respectively, enable a lossless ability to detect gene-based association signals using summary statistics alone.

## 2 Methods

### Algorithm

Our goal is to cluster the variants into groups based on their effects on the multivariate phenotype; that is, we are interested in the joint posterior distribution that indicates cluster memberships per-variant and per-gene. For multi-parameter models such as these, the joint posterior may be difficult to sample from directly. Often, it is easier to sample sequentially from the full conditional distribution of each parameter ^21^ using a Gibbs sampler, a Markov Chain Monte Carlo (MCMC) algorithm that constructs a dependent sequence of parameter values whose distribution approximates the joint posterior ^21–23^. We implement the Gibbs sampler in MRPMM as follows:

- We index genes by *j* = 1, …, *J*, denoting variant *m in gene j* by *v*_*jm*_. Within gene *j*, the variants *v*_*jm*_ are assigned to clusters *c* = 1, …, *C*, each of which is represented by an unscaled effect size parameter ***b***_*c*_.
- The prior for ***b***_*c*_ is N (0, **Θ**_0_), where **Θ**_0_ is an estimate of genetic correlation across the traits.
- Consider the model with *C >* 1 clusters. In order to model the sharing of clusters across the genes, we first draw a *C*-dimensional probability vector ***π***_0_ ∼ Dirichlet (1, 1, 1, …, 1).
- Next, for each gene *j*, we draw a probability vector ***π***_*j*_|***π***_0_ ∼ Dirichlet (*α****π***_0_) to determine the mixture proportions *π*_*jc*_ that dictate how the variants in gene *j* are distributed across the clusters 1, …, *C*.
- The parameter *α* governs how similar the cluster proportions are across genes; it is drawn from a prior *α* ∼ Inv-Gamma (1, 1).
- The algorithm also takes into account the functional annotation, via 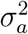, a variance parameter for annotation *a* across the clusters. It follows an inverse-gamma prior 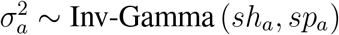, with hyperparameter values *sh*_*a*_ (shape) and *sp*_*a*_ (spread).
- Under the above assumptions of the model, the phenotype of individual *i* with a rare allele of variant *m* with annotation *a* in gene *j* is 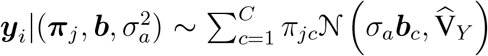, where 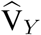 is the estimated residual variance-covariance matrix of phenotypes after the effects of the variants have been regressed out.
- If we have access only to summary statistics of estimated effect sizes 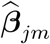 then we estimate variance-covariance matrices 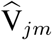 for each variant *m* in gene *j* as 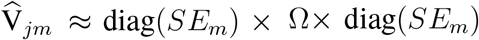, where *SE*_*m*_ denotes the *K*-dimensional vector of standard errors across *K* phenotypes for variant *m*, and Ω is the *K* × *K* matrix of correlation of errors estimated from null variants. Then, our sampling model for the data is 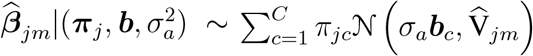.
- Similar to the individual level data model above, this formulation assumes independence between the variants, which is approximately true when each individual carries at most one of the rare variants considered. A connection between the two data types is that 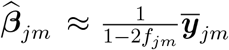, and 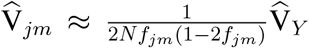, where *f*_*jm*_ is the frequency of the rare allele at variant *m* of gene *j* among the 2*N* haplotypes, and 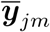 is the mean phenotype of the carrier individuals of that allele.
- To compare models with different numbers of clusters, we use the Bayesian Information Criterion (BIC)^15^, defined for the model M_*C*_ with *C* clusters as

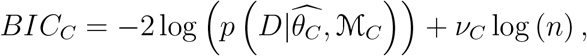

where *D* denotes the observed data, 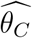 is the vector of maximum likelihood estimates of the parameters of M_*C*_, *ν*_*c*_ is the number of independent parameters of M_*c*_, and *n* is the number of data points. The difference in BIC values between two models approximates twice the logarithm of the Bayes Factor between the models, with lower BIC value corresponding to the model preferred.

MRPMM utilizes Metropolis-Hastings (MH) steps to update the parameters of interest. The algorithm is run for *n*_burn_ + *n*_iter_ iterations, of which the first *n*_burn_ are discarded as an initial “burn-in”. With superscripts in parentheses denoting iteration, the steps of the algorithm are as follows.

1. **Initialize parameters**.
  a. Draw *α*^(0)^ ∼ Inv-Gamma (1, 1).
  b. Draw 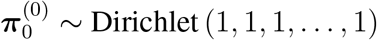.
  c. Draw for each gene 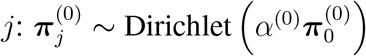;
  d. 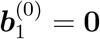.
  e. For each *c* in 2, …, *C* draw 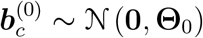.
  f. 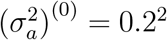 for all *a*.
2. **Repeat the following steps for iterations** *t* = 1, 2, …, *n*_burn_ + *n*_iter_:
  a. Update ***π***_0_ (probability of cluster assignment independent of gene). We use a Metropolis-Hastings sub-step ^24^ using a proposal centred around the current value, drawing 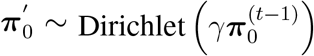. To calculate the acceptance probability, we define the normalizing constant for the C-dimensional Dirichlet distribution with parameters ***z*** as

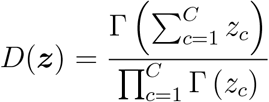

and the density at point ***x*** as

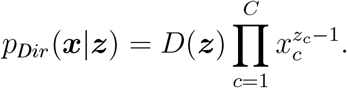

With this notation, the Metropolis-Hastings transition probability from 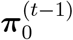to 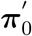 is 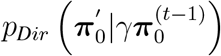, and the density of the observed data depends on 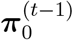 only through the product 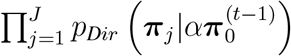. Hence, the proposal acceptance probability is

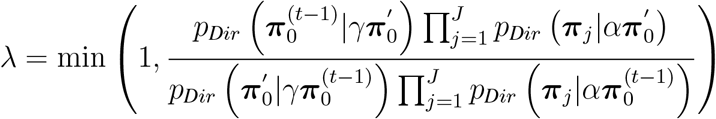

Ergo, with probability *λ*, we set 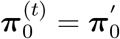, and with probability 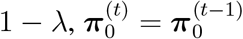.
  b. For each gene *j* = 1, …, *J* we update ***π***_*j*_ (per-gene cluster parameter) to be

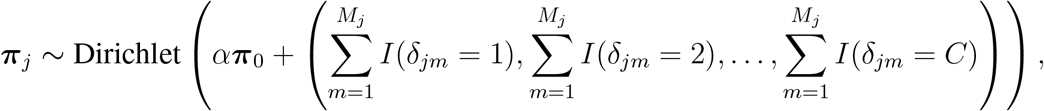

where *δ*_*jm*_ is the index of the cluster to which the variant *m* of gene *j* belongs to and *I*(·) is the indicator function.
  c. Update *δ*_*jm*_ for all *j* = 1, …, *J* and *m* = 1, …, *M*_*j*_. For all *c*, compute

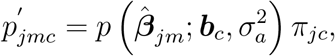

and renormalize in such a way that 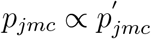 sums to 1 over all *c*. Then, we sample

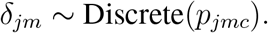
  d. Update ***b***_*c*_ (cluster effect profile) using a Gibbs update from a Gaussian distribution:

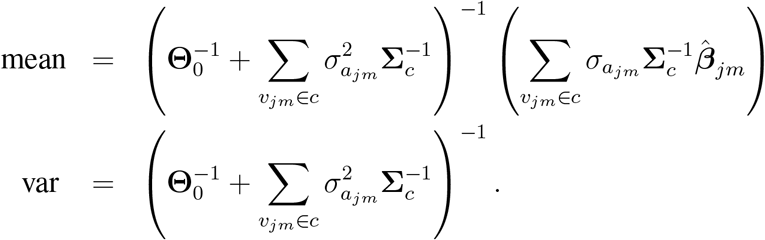
  e. Update 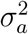 (annotation spread parameter). We use a Metropolis-Hastings sub-step using a random-walk proposal. That is, sequentially for each annotation *a*, we sample a proposal value

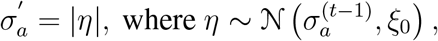

where *ξ*_0_ is a hyperparameter controlling the spread of the proposals. In the examples, we have used *ξ*_0_ = 1. Then we calculate the acceptance probability

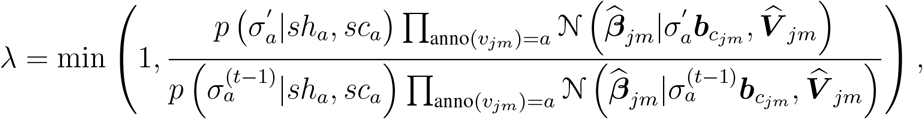

where the products are over those variants *v*_*jm*_ whose annotation is *a* and *c*_*jm*_ is the cluster of variant *v*_*jm*_. With probability *λ*, we set 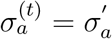, and with probability 1 − *λ*, 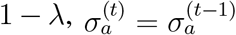
  f. Update *α* (parameter determining sharing of clusters across genes). We again use a Metropolis-Hastings sub-step using a random-walk proposal. That is, we sample a proposal value *α*’
s = |*η*| where *η* ∼ 𝒩(*α*^(*t*−1)^, *ξ*_*α*_) where *ξ*_*α*_ is a fixed value controlling the variance of the proposal distribution. Then, we compute the acceptance probability

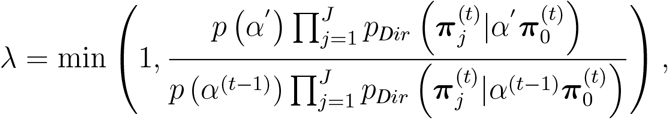

where the prior for *α* is Inv-Gamma(1,1) (i.e., *p*(*α*) = *α*^−2^ exp (−*α*^−1^)), and *p*_*Dir*_ is the density of the Dirichlet distribution as defined above in part (a). With probability *λ*, we set *α*^(*t*)^ = *α*, and with probability 1 − *λ, α*^(*t*)^ = *α*^(*t*−1)^.

### Data

We used a combination of self-reported ancestry (UK Biobank field ID 21000), principal component analysis on genotype data, and the relatedness matrix to identify six subpopulations in the study: white British, African, South Asian, non-British white, semi-related, and an admixed population. To determine the first four populations, which contain samples not related closer than the third degree, we first used the principal components of the genotyped variants from the UK Biobank and defined thresholds on principal component 1 and principal component 2 and further refined the population definition as described elsewhere^25^ Semi-related individuals were grouped as individuals whose genetic data (after passing UK Biobank QC filters; sufficiently low missingness rates; and genetically inferred sex matching reported sex), using a King’s relationship table, were between conditional third and conditional second degrees of relatedness. Admixed individuals were grouped as unrelated individuals who were flagged as “used_in_pca_calculation” by the UK Biobank and were not assigned to any of the other populations^6^.

We performed genome-wide association analysis on individuals with whole-exome sequencing data for three lipid-related phenotypes (high-density lipoproteins [UK Biobank Field 30760], triglycerides [UK Biobank Field 30870], and low-density lipoproteins [UK Biobank Field 30780]) and three renal-related phenotypes (creatinine [UK Biobank Field 30700], cystatin C [UK Biobank Field 30720], and effective glomerular filtration rate [derived from UK Biobank Field 30700]). The analyses were performed for each of the six population subgroups as defined above using PLINK v2.00a (20 October 2020). The quantitative trait values were rank normalized using the – pheno-quantile-normalize flag. We used age, sex, and the first ten genetic principal components as covariates in the analyses. The analysis was performed for 5,850,789 rare (minor allele frequency ≤ 0.01) protein-truncating (492,151) and protein-altering (5,358,638) variants.

For the admixed population, we conducted local ancestry-corrected GWAS. We first assembled a reference panel from 1,380 single-ancestry samples in the 1000 Genomes Project^26^, the Human Genome Diversity Project^27^ and the Simons Genome Diversity Project^28^ choosing appropriate ancestry clusters by running ADMIXTURE^29^ with the unsupervised setting. Using cross-validation, eight well-supported ancestral population clusters were identified: African, African Hunter-Gatherer, East Asian, European, Native American, Oceanian, South Asian, and West Asian. We then used RFMix v2.03^30^ to assign each of the 20,727 windows across the phased genomes to one of these eight ancestry clusters (for all individuals in the UK Biobank). These local ancestry assignments were subsequently used with PLINK2 as local covariates in the GWAS for the admixed individuals for SNPs within those respective windows. PLINK2 allows for the direct input of the RFMix output (the MSP file, which contains the most likely subpopulation assignment per conditional random field [CRF] point) as local covariates using the “local-cov”, “local-psam”, and “local-haps” flags, the “local-cats0=n” flag (where n is the number of assignments), and the “local-pos-cols=2,1,2,7” flag (for a typical RFMix MSP output file - see “Association Analysis” page on PLINK website).

Subsequently, we used METAL^31^ to perform inverse-variance weighted meta-analysis to generate a single summary statistic file per phenotype.

For the remainder, we used Variant Effect Predictor (VEP)^32^ to annotate the most severe consequence, the gene symbol, and HGVSp of each variant in the UK Biobank exome data. We calculated minor allele frequencies using PLINK. We provide these metadata, which are necessary for MRPMM, in exome and array tables, available for direct download via the Global Biobank Engine^33^ (Code Availability).

## 3 Results

To assess MRPMM’s ability to estimate the underlying mixture of effects from summary statistics, we chose two sets of related phenotypes: one set of lipid phenotypes (high-density lipoprotein cholesterol levels [HDL], low-density lipoprotein cholesterol levels [LDL], and triglycerides [TG]), and another set of renal phenotypes (creatinine [CRE], cystatin C [CSTC] and effective glomerular filtration rate [eGFR]). We then identified genes with a significant burden of associated, rare, protein-truncating variants (PTVs) in a meta-analysis comprising 184,698 UK Biobank individuals amongst six cohorts of different ancestries (white British [137,920], non-British white [10,432], African [2,716], South Asian [3,569], semi-related [18,100], and admixed [11,961]). After using a Bayesian model comparison approach^6^ to consider GWAS evidence across each set of phenotypes and across each gene, we chose 13 genes that had a log_10_ Bayes Factor (BF) ≥ 5 for lipid traits, and 5 genes that met that criterion for renal traits. We then ran MRPMM across PTVs in these genes, increasing the number of hypothesized clusters until there was an increase in BIC, a goodness-of-fit measure which is minimized under ideal conditions. We found that four clusters (including the “null cluster” with a constant effect of 0 across phenotypes) was favored for both sets of phenotypes (Figure 1), and that each cluster had a distinct effect size profile on the multivariate phenotype (Figures 2, 3).

**Figure 1a:**
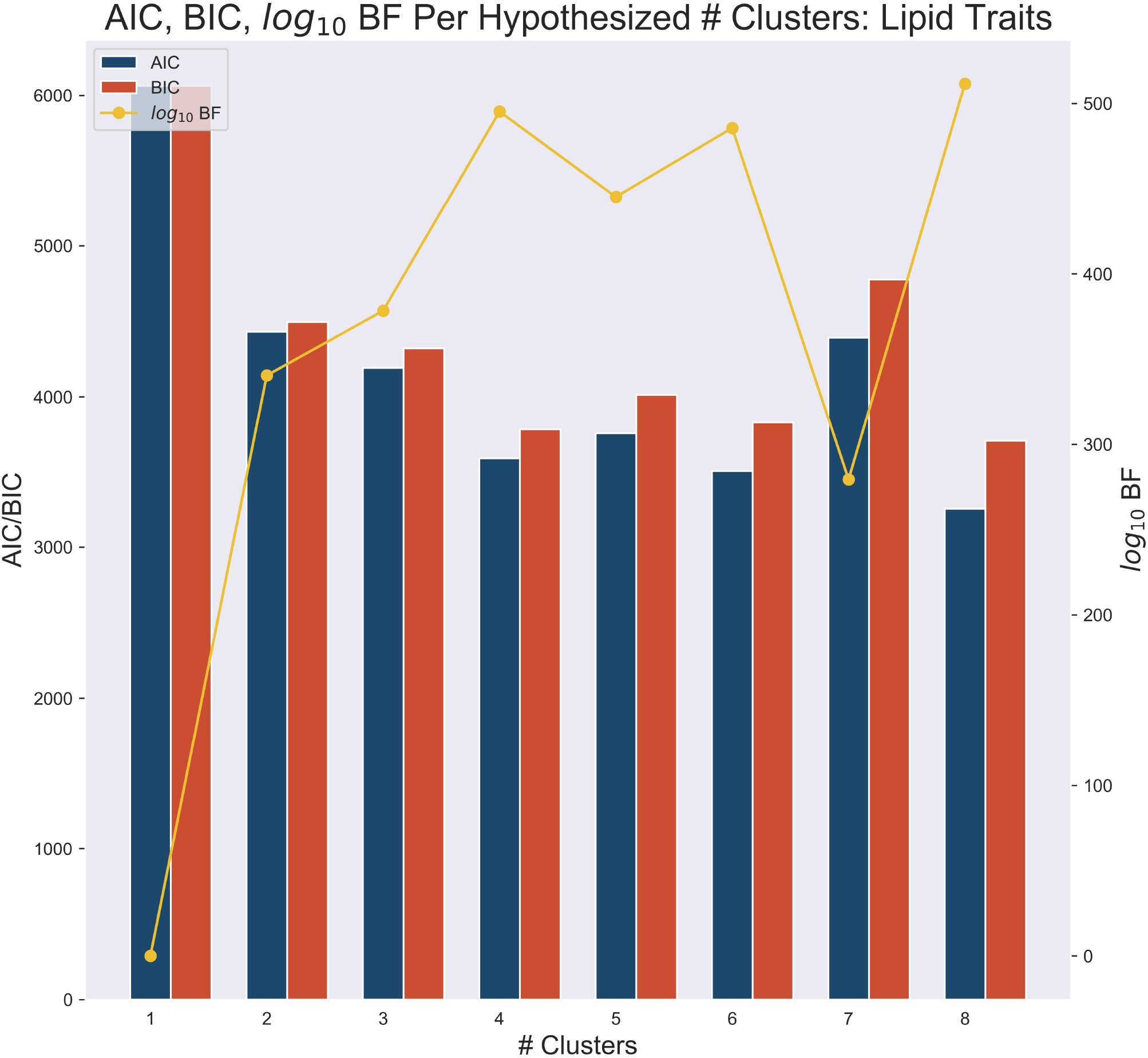
AIC, BIC (goodness-of-fit), and log_10_ BF as compared to the null cluster for number of clusters of effects on a lipid-related multivariate phenotype of high-density lipoprotein cholesterol (HDL), triglyceride (TG) levels, and low-density lipoprotein cholesterol (LDL). The algorithm was stopped when these AIC or BIC values trended upwards and log_10_ BF was maximized. In this case, we stopped at 5 and chose 4 clusters.

**Figure 1b:**
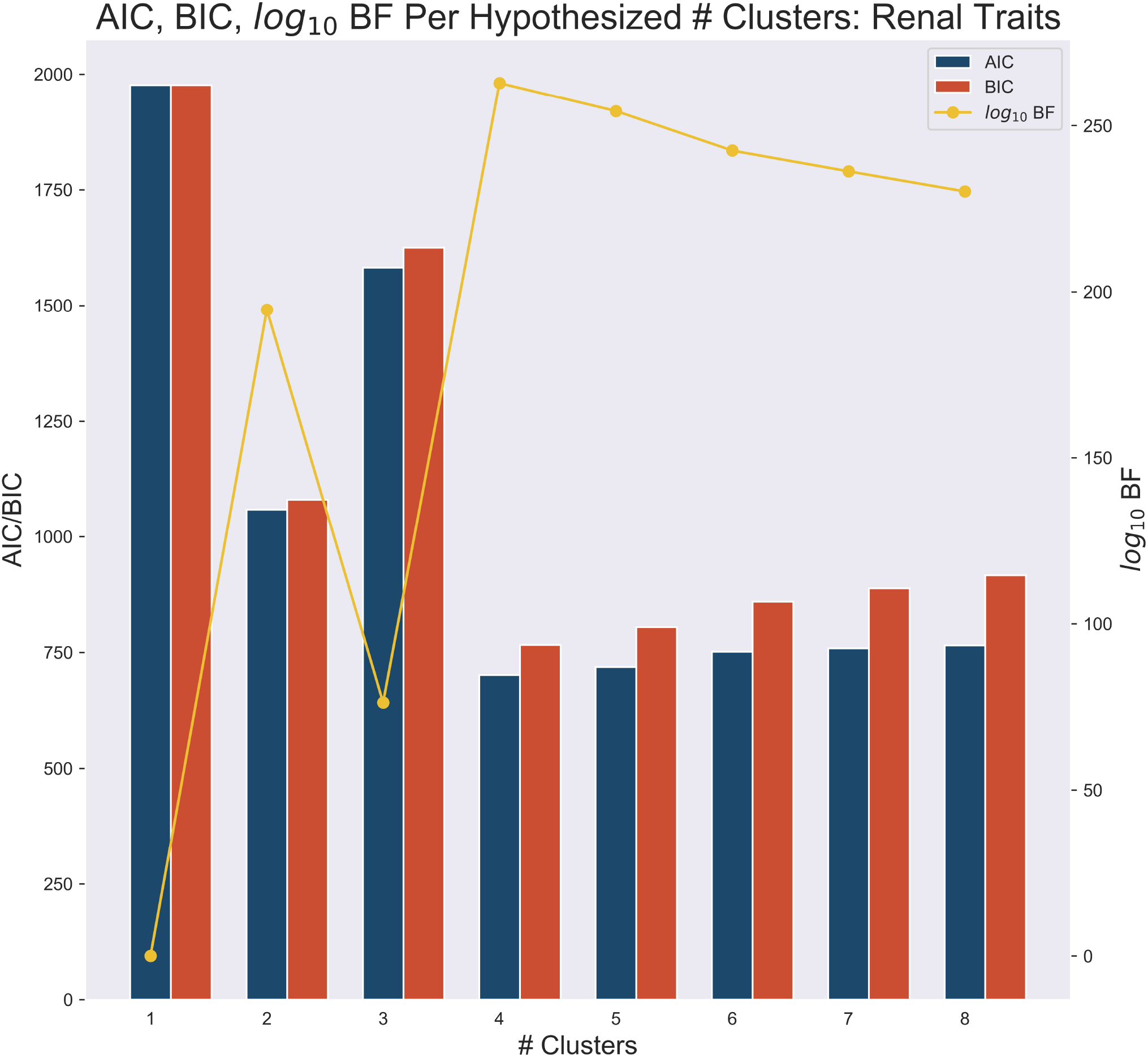
AIC, BIC (goodness-of-fit), and log_10_ BF as compared to the null cluster for number of clusters of effects on a renal-related multivariate phenotype of creatinine (CRE), cystatin C (CSTC) levels, and effective glomerular filtration rate (eGFR). The algorithm was stopped when these AIC or BIC values trended upwards and log_10_ BF was maximized. In this case, we stopped at 5 and chose 4 clusters.

**Figure 2:**
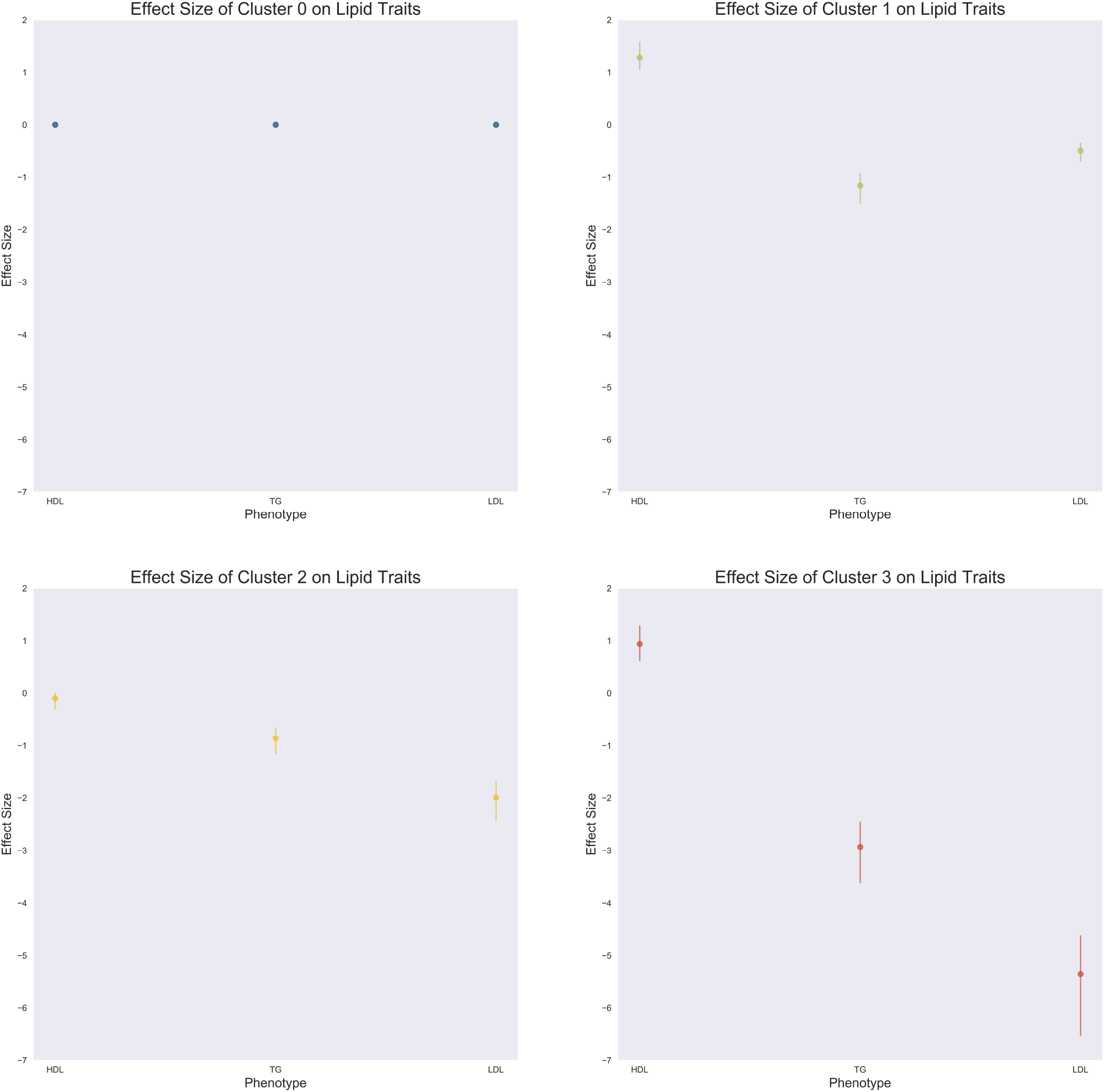
**a (upper left):** Effect sizes of Cluster 0 (with 95% credible intervals) on the lipid-related multivariate phenotype (the “null cluster”). **b (upper right):** Effect sizes of Cluster 1 (with 95% credible intervals) on the lipid-related multivariate phenotype. PTVs in this cluster espouse positive effects on HDL levels and negative effects on TG levels and LDL levels. **c (lower left):** Effect sizes of Cluster 2 (with 95% credible intervals) on the lipid-related multivariate phenotype. PTVs in this cluster espouse null effects on the HDL phenotype and successively negative effects on TG and LDL. **d (lower right):** Effect sizes of Cluster 3 (with 95% credible intervals) on the lipid-related multivariate phenotype. This cluster seems to have disparate effects on HDL (positive effect) as compared to TG (strong negative effect) and LDL (extremely strong negative effect).

**Figure 3:**
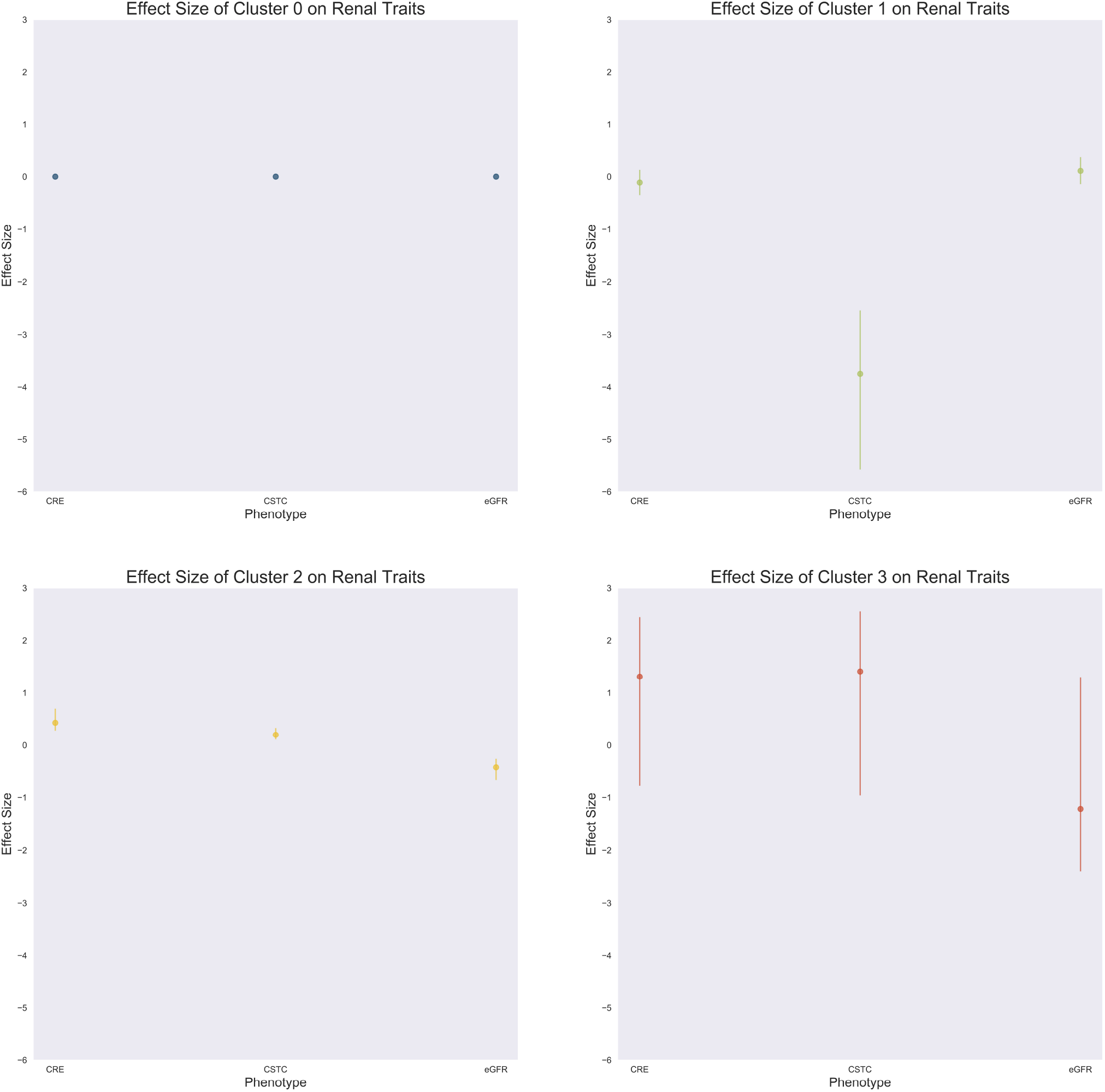
**a (upper left):** Effect sizes of Cluster 0 (with 95% credible intervals) on the renal-related multivariate phenotype (the “null cluster”). **b (upper right):** Effect sizes of Cluster 1 (with 95% credible intervals) on the renal-related multivariate phenotype. PTVs in this cluster espouse null effects on CRE and eGFR levels and strong negative effects on CSTC levels. **c (lower left):** Effect sizes of Cluster 2 (with 95% credible intervals) on the renal-related multivariate phenotype. PTVs in this cluster espouse mild positive effects on CRE and CSTC and mild negative effects on eGFR. **d (lower right):** Effect sizes of Cluster 3 (with 95% credible intervals) on the renal-related multivariate phenotype. This cluster seems to have inconclusive effects on the multivariate phenotype, as the 95% credible intervals for the effects cross 0.

Specifically, we see that there are some *APOB* variants (Figure 4a - left side) that have a strong negative effect on TG and LDL levels (Figure 2d) through exclusive membership to Cluster To the right of those variants, other variants are definitively placed into Cluster 2 and Cluster 1 respectively. The variants that are definitively placed into Cluster 2 feature the *PCSK9* and *ANGPTL3* genes; MRPMM shows that these variants down-regulate TG and LDL levels (Figure 2c). The variants which belong exclusively to Cluster 1 feature the *PDE3B, APOC3, CETP*, and *ANGPTL8* genes and have positive effects on HDL levels, negative effects on TG levels, and mild negative effects on LDL levels (Figure 2b). We can perform a similar visual analysis with the renal multivariate phenotype results (Figure 4b). Variants from the *CGNL1, RNF186, SLC22A2*, and *SLC34A3* genes largely seem to fall under Cluster 2, whereas variants from *CST3* clearly fall into Cluster 1 in an isolated manner. Cluster 3 seems to be populated partially by several variants in *CGNL1*. These variant-specific breakdowns and their gene-aggregate counterparts (Figures 5a, 5b) help trace back the variants’ effect profile on the multivariate phenotype. Unlike in aggregation approaches, where a single statistic captures which genes are associated with the trait without providing much context as to the nature of the association, MRPMM has the useful characteristic of being able to cluster not only variants but also genes into different effect profiles. We provide single-trait MRPMM results for all traits across the UK Biobank on the Global Biobank Engine^33^ (Code Availability).

**Figure 4a:**
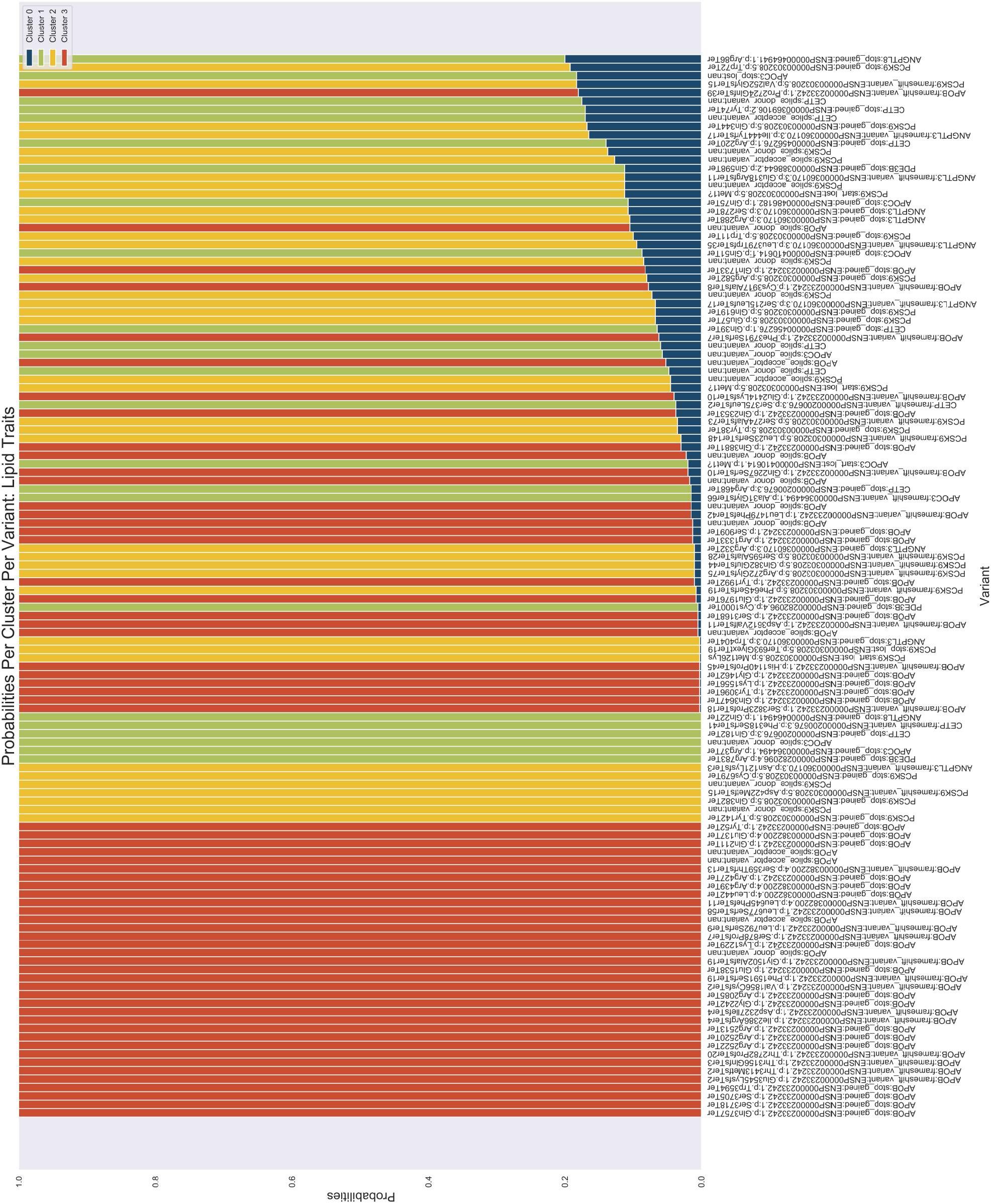
Variant-level posterior probabilities that PTVs in candidate genes in the UK Biobank exome sequencing data set belong to each of four clusters hypothesized with respect to a lipid-related multivariate phenotype of high-density lipoprotein cholesterol (HDL), low-density lipoprotein cholesterol (LDL), and triglyceride (TG) levels. The PTVs in *APOB* that are marked as belonging to Cluster 3 on the left are responsible for coding for a hepatic lipase enzyme, hence their strong effects on triglycerides and LDL cholesterol levels (see Figure 3D). Only PTVs that have posterior probability ≤0.2 of belonging in the null cluster (Cluster 0) are shown. Variants are sorted by the posterior probability of belonging to the null cluster.

**Figure 4b:**
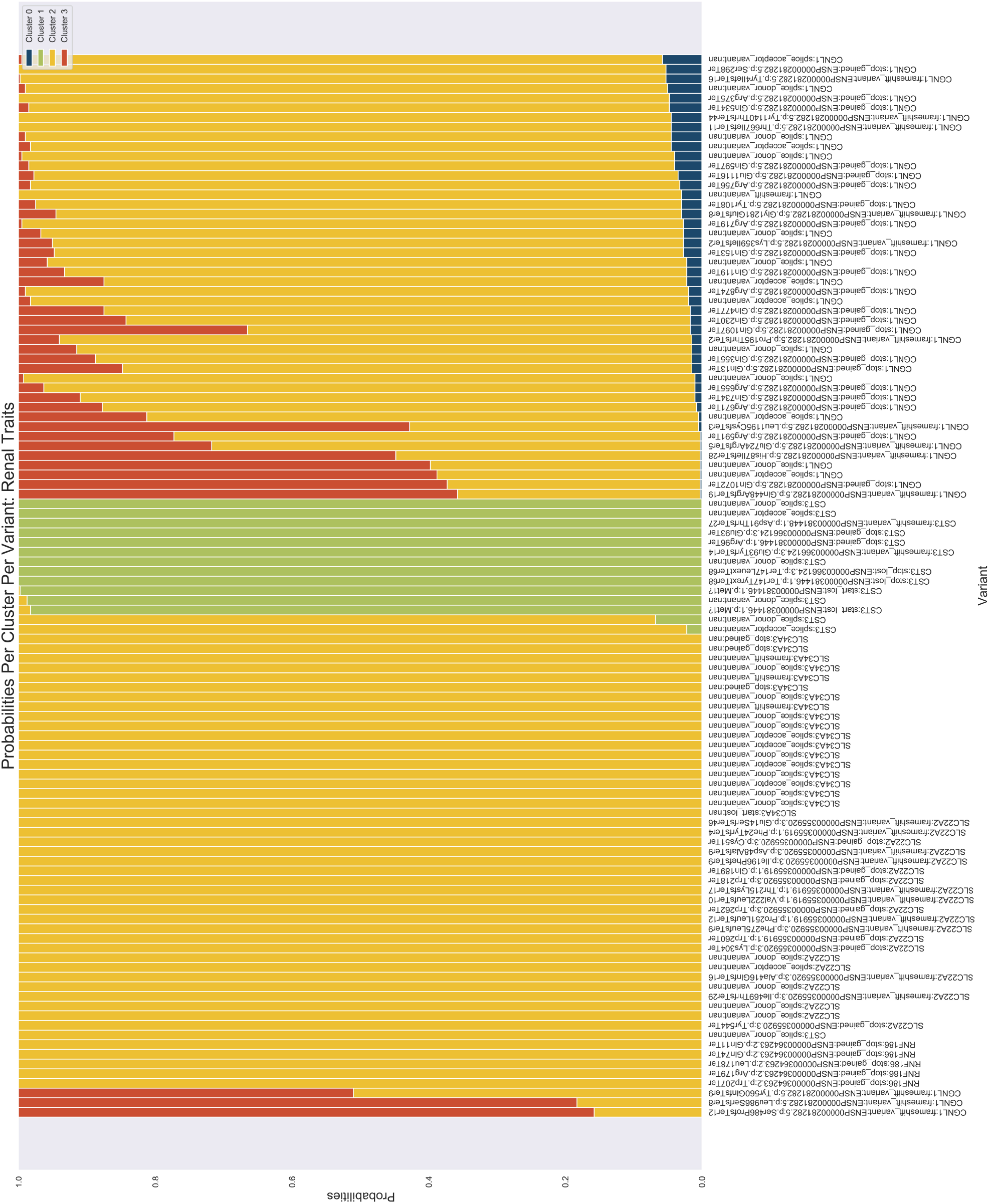
Variant-level posterior probabilities that PTVs in candidate genes in the UK Biobank exome sequencing data set belong to each of four clusters hypothesized with respect to a renal-related multivariate phenotype of creatinine (CRE), cystatin C (CSTC), and effective glomerular filtration rate (eGFR). The PTVs in *CST3* that are marked as belonging to Cluster 1 in the middle are responsible for coding cystatin C directly, hence their strong effects CSTC levels (see Figure 4B). Only PTVs that have posterior probability ≤0.2 of belonging in the null cluster (Cluster 0) are shown. Variants are sorted by the posterior probability of belonging to the null cluster.

**Figure 5a:**
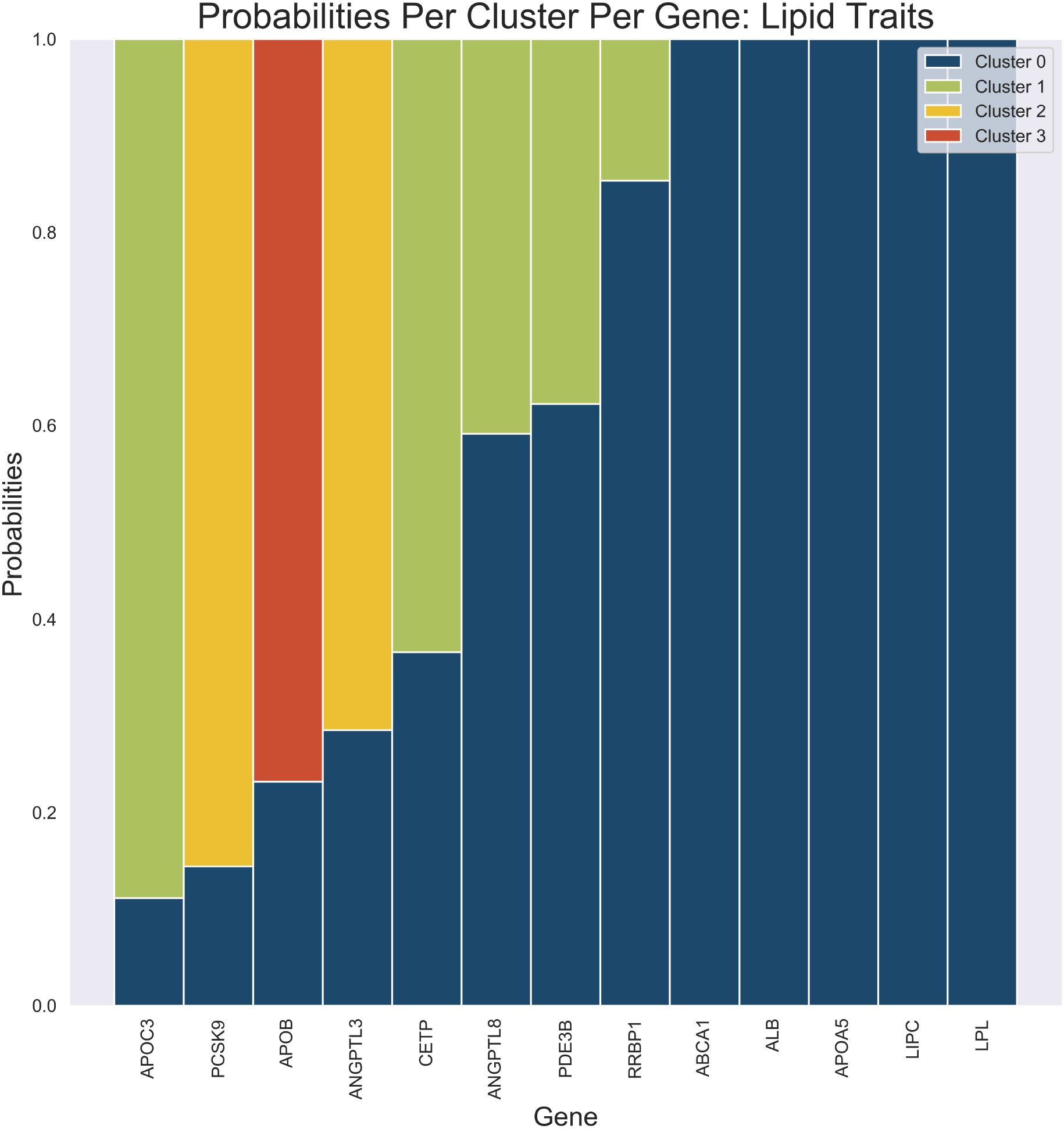
Gene-level posterior probabilities that PTVs in candidate genes in the UK Biobank exome sequencing data set belong to each of four clusters hypothesized with respect to a lipid-related multivariate phenotype of high-density lipoprotein cholesterol (HDL), low-density lipoprotein cholesterol (LDL), and triglyceride (TG) levels. All PTVs in the analysis are accounted for here.

**Figure 5b:**
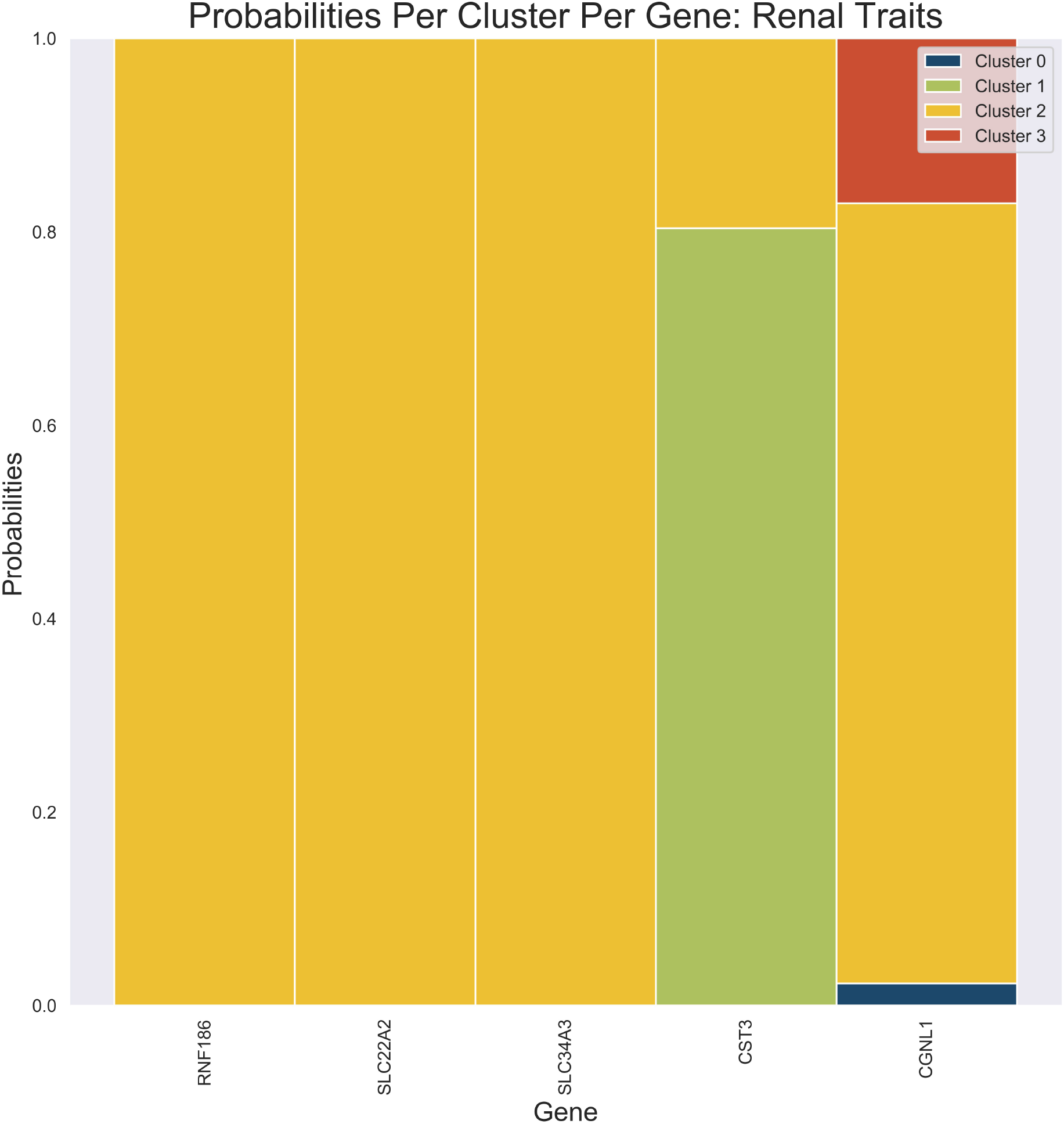
Gene-level posterior probabilities that PTVs in candidate genes in the UK Biobank exome sequencing data set belong to each of four clusters hypothesized with respect to a renal-related multivariate phenotype of creatinine (CRE), cystatin C (CSTC), and effective glomerular filtration rate (eGFR). All PTVs in the analysis are accounted for here.

## 4 Future Directions and Discussion

In this study, we used a Bayesian hierarchical mixture model to estimate the underlying mixture of components driving association signals between various protein-truncating variants and a set of genetically related phenotypes. By explicitly modeling the sharing of effects across genes, we are able to use Gibbs sampling to approximately infer the joint posterior distribution and thereby assign variants to clusters. In both applications, we see that clusters have vastly different effect size profiles on the sets of phenotypes chosen; this shows that aggregating rare-variant signal in blocks (e.g. genes) may not fully encapsulate the information that is available in summary statistics. Coupling a Bayesian model comparison approach as described by Venkataraman et al.^6^ with MRPMM may be a way to (1) systematically screen for genes associated with the set of phenotypes of interest and then (2) cluster the effect size profiles of rare variants within these genes, thereby providing a window into the underlying biology. For example, aligning the results from the MRPMM analysis and other function-elucidating analyses like protein domain models or 3D structure analysis could potentially lead to the identification of promising therapeutic targets. Going forward, it is also essential that this analysis be performed using whole genome sequencing data; as MRPMM is able to use any type of annotation, how variant effects translate across epigenomic profiles and/or conservation patterns may become relevant and useful to analyze in these settings. Overall, MRPMM provides interpretability at the level of individual variants, in contrast to typical rare-variant techniques that work only at the level of aggregated variants.

## 5 Code Availability

We have published the full set of associations (log_10_ BF ≥ 5) from an independent effects model amongst PAVs, from a similar effects model amongst PAVs, as well as from a similar effects model amongst PTVs on the Global Biobank Engine^33^. While this study focuses on exome associations (https://biobankengine.stanford.edu/RIVAS_HG38/mrpgene/all), we also provide associations for array data (https://biobankengine.stanford.edu/RIVAS_HG19/mrpgene/all). For every phenotype, we provide single-phenotype MRPMM results (right-most columns in the tables) for PAVs and PTVs, with cluster effect size estimates as well as cluster assignment probabilities and proportions displayed.

Exome and array metadata tables are available on the Global Biobank Engine for direct download at these links:

https://biobankengine.stanford.edu/static/ukb_exm_oqfe-consequence_wb_maf_gene_ld_indep_mpc_pli.tsv.gz - Exome

https://biobankengine.stanford.edu/static/ukb_cal-consequence_wb_maf_gene_ld_indep_mpc_pli.tsv.gz - Array

MRPMM was implemented using Python (dependencies: pandas v1.1.5, numpy v1.16.4, sklearn 0.24.0, scipy v1.3.0). The requirements, code, example usages, and interpretation of results files can be found at https://github.com/rivas-lab/mrpmm.

## 6 Acknowledgments

This research was conducted using the UK Biobank Resource under application number 24983, ‘Generating effective therapeutic hypotheses from genomic and hospital linkage data’ (http://www.ukbiobank.ac.uk/wp-content/uploads/2017/06/24983-Dr-Manuel-Rivas.pdf). Based on the information provided in protocol 44532, the Stanford IRB has determined that the research does not involve human subjects as defined in 45 CFR 46.102(f) or 21 CFR 50.3(g). All participants in the UK Biobank study provided written informed consent (more information is available at https://www.ukbiobank.ac.uk/2018/02/gdpr/). Statin adjustment analyses were further conducted via UK Biobank application 7089 using a protocol approved by the Partners HealthCare Institutional Review Board. We thank all the participants in the UK Biobank. We thank members of the Rivas lab for their feedback. The content is solely the responsibility of the authors and does not necessarily represent the official views of the NIH. Some of the computing for this project was performed on the Sherlock cluster at Stanford University. We would like to thank Stanford University and the Stanford Research Computing Center for providing computational resources and support that contributed to these research results. G.R.V. is supported by the National Library of Medicine (NLM) T15 Continuing Education Training Grant. Y.T. is supported by a Funai Overseas Scholarship from the Funai Foundation for Information Technology and the Stanford University School of Medicine. M.A.R. is in part supported by the NHGRI of the NIH under award R01HG010140 (M.A.R.) and an NIH Center for Multi- and Trans-ethnic Mapping of Mendelian and Complex Diseases grant (5U01 HG009080).

